# Towards risk-based surveillance of African Swine Fever in Switzerland

**DOI:** 10.1101/2021.05.17.444420

**Authors:** Maria Elena Vargas Amado, Luís Pedro Carmo, John Berezowski, Claude Fischer, Maria João Santos, Rolf Grütter

## Abstract

African Swine Fever (ASF) has emerged as a disease of great concern to swine producers and government disease control agencies because of its severe consequences to animal health and the pig industry. Early detection of an ASF introduction is considered essential for reducing the harm caused by the disease. Risk-based surveillance approaches have been used as enhancements to early disease epidemic detection systems in livestock populations. Such approaches may consider the role wildlife plays in hosting and transmitting a disease. In this study, a novel method is presented to estimate and map the risk of introducing ASF into the domestic pig population through wild boar intermediate hosts. It makes use of data about hunted wild boar, rest areas along motorways connecting ASF affected countries to Switzerland, outdoor piggeries, and forest cover. These data were used to compute relative wild boar abundance as well as to estimate the risk of both disease introduction into the wild boar population and disease transmission to domestic pigs. The way relative wild boar abundance was calculated adds to the current state of the art by considering the effect of beech mast on hunting success and the probability of wild boar occurrence when distributing relative abundance values among individual grid cells. The risk of ASF introduction into the domestic pig population by wild boar was highest near the borders of France, Germany, and Italy. On the north side of the Alps, areas of high risk were located on the unshielded side of the main motorway crossing the Central Plateau, which acts as a barrier for wild boar. The results of this study can be used to focus surveillance efforts for early disease detection on high risk areas. The developed method may also inform policies to control other diseases that are transmitted by direct contact from wild boar to domestic pigs.

## Introduction

Wild boar represent a health threat to domestic pigs (Laddomada, et al. 1994, Fritzemeier, et al. 2000, Köppel, et al. 2007, Ruiz-Fons, et al. 2008, Wu, Abril and Hinicacute, et al. 2011), because they share susceptibility to a range of infectious diseases with domestic breeds. Diseases found in wild boar that are a significant threat to the swine industry include: classical swine fever, Aujeszky’s disease, and porcine brucellosis (Köppel, et al. 2007, Ruiz-Fons, et al. 2008). In Switzerland, not only has the wild boar population increased in the last decades (Sáez-Royuela and Tellería 1986, Geisser and Reyer 2004, Massei, et al. 2015), but the number of outdoor piggeries has also grown. With these two developments, the probability of contact between free ranging wild boar and farmed pigs has increased (Köppel, et al. 2007). Recently African Swine Fever (ASF) has emerged as a disease of great concern to swine producers and government disease control agencies because of its health and economic consequences. It re-emerged in Eurasia in 2007 (Vergne, Gogin and Pfeiffer 2017), jumping to East Europe in 2014 (Gallardo, et al. 2018), to Belgium in 2018 (Morelle, et al. 2019), and more recently, in 2020, the first case was reported in Germany (Landwirtschaftsverlag 2020).

In Switzerland, domestic pigs have a relatively high health status and are free from many diseases including ASF (Köppel, et al. 2007, Nathues, et al. 2016). However, ASF outbreaks have occurred quite close to the Swiss borders and ASF poses a substantial threat with potentially extreme consequences to the Swiss pig industry. Early detection of an ASF introduction will be essential for reducing the harm caused by the disease. Risk-based surveillance approaches have been widely used as enhancements to early epidemic detection systems in livestock populations. For instance, in Great Britain risk-based approaches were used to identify high risk areas where surveillance should be focused in order to detect avian influenza outbreaks (Snow, et al. 2007). In New Zealand, risk-based surveillance was used to detect vector-borne diseases causing ovine and caprine abortion (Prattley 2009). A risk assessment framework was used to determine the probability of infection of European swine with the ASF virus through wild boar movement and legal trade of pigs and pig meat (Taylor, et al. 2020). Risk assessment in that study was performed at a fine spatial scale, allowing the limited surveillance and intervention resources to be focused on high-risk areas and pathways. In Switzerland, the benefits of implementing risk-based surveillance approaches have been reported for 1) freedom from infectious bovine rhinotracheitis (IBR) and enzootic bovine leucosis (EBL), 2) bluetongue surveillance, and 3) the national residue monitoring program (Reist, Jemmi and Stärk 2012).

In order to assess the risk of occurrence of an ASF outbreak within the wild boar population in Switzerland, it is important to know the spatial distribution and relative abundance of wild boar. Density and abundance estimations are widely used to monitor, manage, and control wildlife populations (Pittiglio, Khomenko and Beltran-Alcrudo 2018). This information is used by authorities (Acevedo, et al. 2007) to assess the vulnerability of crops to damage by wild boar (Geisser and Reyer 2004, Honda and Kawauchi 2011) or to implement population control activities such as fencing, trapping, and hunting (Chapman and Trani 2007).

Information about potential transmission routes is also needed as it can be used to focus wild boar ASF surveillance activities on geographical areas where there is a high risk of pathogen introduction. One way of introducing the disease is by improper disposal of contaminated food waste in areas where wild boars are known to be present (Mur, et al. 2012, EFSA 2010). This was suspected in Belgium in 2018 (Belgian Federal Agency for the Safety of the Food Chain (FASFC) 2019). Travelers coming from countries where the disease is currently active can introduce the pathogen through contaminated food that is disposed of in rest areas along motorways. Wild boar are opportunistic scavengers (Penrith and Vosloo 2009), and if discarded food is improperly contained, they may consume it and become infected, providing pathway for the pathogen to enter the wild boar population.

If there is an introduction of ASF into the Swiss wild boar population, it is likely that the initial spread of the pathogen will occur locally among wild boar. Because the pathogen can be easily transmitted by direct contact from wild boar to domestic pigs, it is important to identify pig holdings in close proximity to wild boar populations where a direct contact could potentially occur. Knowing the location of these holdings is essential for optimizing surveillance for early detection of an ASF introduction into domestic swine. Once the pathogen is introduced into the domestic pig population, initial spread is also likely to be local. The most rapid spread of the introduced pathogen is expected in geographical areas with the greatest density of the most highly connected piggeries. Because of the severe consequence of an ASF introduction into these areas, they should also be a focus for early epidemic detection surveillance. Once the disease enters one node (farm) of the pig production network, the spread across the entire pig production network can potentially be very fast, compromising the swine production supply chain and Swiss export markets for pigs and pig products (Stärk, et al. 2006).

This study provides information to support the implementation of a risk-based surveillance system for ASF entering Switzerland by contaminated food waste, including 1) identifying risk areas that could represent entrance points of ASF into the wild boar population by identifying geographic areas where there are high relative abundances of wild boar and rest areas along important motorways, 2) identifying the outdoor piggeries in which domestic pigs may be more likely to be exposed to the ASF virus due to a high relative abundance of wild boar, and 3) identifying areas with a combined risk of introducing ASF into the domestic pig population by wild boar.

In a previous study, the potential distribution of wild boar in Switzerland was modeled (Vargas-Amado, et al. 2020). In the current study we enhanced this information by modeling the effect of beach mast on hunting success in order to calculate wild boar relative abundance in Switzerland, with a fine-grained spatial resolution using hunting statistics as input data.

## Material and methods

### Study area

The study considers all of Switzerland, a country that covers a total surface area of 41,285 sq km ranging from 193 to 4,634 meters above sea level (Swiss Confederation 2020a). Seven and a half percent of Switzerland’s surface area is used for settlements and urban areas, trade, industry and transport, energy supply and waste disposal or recreational areas and parks. Agricultural land occupies 35.9%, and forests and woodlands 31.3%. Switzerland has three main geographic regions: the Alps, covering around 60% of the country’s total surface area, the Swiss Plateau (30%) and the Jura (10%). The Alps act as a prominent climatic barrier between Northern and Southern Switzerland (Swiss Confederation 2020b). The climate of Northern Switzerland is heavily influenced by the Atlantic Ocean. Winters in the Northern Plateau are mild and damp, whereas higher altitudes experience arctic temperatures. At altitudes above 1,200– 1,500 meters, precipitation in winter mainly falls as snow. Southern Switzerland is strongly affected by the Mediterranean Sea, making winters mild and summers warm and humid, and sometimes hot.

### Data collection

#### Hunting data

Hunting data from 2011/12–2017/18 were the primary data source for the computation of relative wild boar abundance. They were obtained from all cantons in which, according to the Federal Hunting Statistics, wild boar are present, except Basel-Stadt and Luzern. For the latter two cantons, the data reported in the Federal Hunting Statistics were used. The data from Vaud were obtained only for the period from 2012/13–2017/18, those from Fribourg were obtained for the period from 2013/14– 2017/18. These longitudinal data made it possible to balance out the strong effects of non-controllable factors on the annual number of harvested wild boar. For instance, weather conditions such as snow cover and snow depth strongly influence the efficiency of hunting by making some areas less accessible to hunters (ENETWILD-consortium, et al. 2018). The aggregate data used in this study are reported per canton and year in the Federal Hunting Statistics.^1^ Both the spatial and the temporal granularity of the data varied widely between different cantons, ranging from daily data with exact geographic location (i.e., coordinates) to yearly data aggregated per canton (see Table 1). This heterogeneity required several preprocessing steps in order to make the data comparable before computing relative abundance (see Section ‘Computation of relative wild boar abundance’).

**Table 1.** The 26 cantons of Switzerland categorized according to the temporal and spatial granularity of the available hunting data. Category 0 represents cantons where wild boar, according to the hunting authorities, are not yet present. Some communes (value ‘Comm’) or hunting grounds (value ‘Rev’) in categories 2–4 were subject to mergers during the observation period and required particular attention. The canton of Geneva is a special case, because hunting is prohibited throughout the entire year (still between 150 and 200 wild boar are shot every year).

| No. | Name | Code | Temporal | Spatial | Hunting Season |
| --- | --- | --- | --- | --- | --- |
| 0 | Schwyz | SZ | N/A | N/A | N/A |
| 0 | Obwalden | OW | N/A | N/A | 01-09 to 28-02 |
| 0 | Glarus | GL | N/A | N/A | 01-09 to 30-11 |
| 0 | Uri | UR | N/A | N/A | 01-09 to 31-12 |
| 0 | Zug | ZG | N/A | N/A | 01-10 to 31-01 |
| 0 | Nidwalden | NW | N/A | N/A | 01-07 to 28-02 |
| 1 | Appenzell Innerrhoden | AI | Day | Coord | 04-09 to 31-01 |
| 1 | Neuchâtel | NE | Day | Coord | 13-08 to 28-02 |
| 1 | Vaud | VD | Day | Coord | 01-06 to 09-02 |
| 1 | Graubünden | GR | Day | Coord | 01-09 to 20-12 |
| 1 | Fribourg | FR | Day | Coord | 01-07 to 31-01 |
| 1 | Zurich | ZH | Day | Coord | 01-07 to 28-02 |
| 1 | St. Gallen | SG | Day | Coord | 01-07 to 28-02 |
| 2 | Appenzell Ausserrhoden | AR | Day | Comm | 01-08 to 31-01 |
| 2 | Ticino | TI | Day | Comm | 01-09 to 31-01 |
| 2 | Valais | VS | Day | Comm | 17-09 to 27-01 |
| 3 | Jura | JU | Day | Comm | 15-06 to 28-02 |
| 2 | Aargau | AG | Day | Rev | 01-07 to 31-01 |
| 3 | Bern | BE | Day | Comm/Coord | 02-08 to 31-01 |
| 3 | Solothurn | SO | Day | Rev/Coord | 01-07 to 28-02 |
| 4 | Basel-Landschaft | BL | Year | Comm | 01-07 to 28-02 |
| 4 | Schaffhausen | SH | Year | Rev | 01-07 to 28-02 |
| 4 | Thurgau | TG | Year | Rev | 01-07 to 28-02 |
| 5 | Basel-Stadt | BS | Year | Canton | 01-07 to 28-02 |
| 5 | Luzern | LU | Year | Canton | 01-07 to 28-02 |
| N/A | Geneva | GE | N/A | N/A | N/A |

#### Hunting calendar

The calendar days falling within the hunting period were extracted from the Federal Hunting Statistics for each canton (Table 1). They were used to compute the hunting effort on as granular a spatial level as possible (see Section ‘Computation of relative wild boar abundance’).

#### Beech mast index

Available food resources, among them the fruit of forest trees, have a strong influence on winter survival and spring reproduction of wild boar (Frauendorf, et al. 2016, Gamelon, et al. 2017, Geisser and Reyer 2005, Vetter, et al. 2015). Fruit production of tree species such as beech varies from year to year. Years with high fruit production are called mast years. Based on phenomenological criteria a four-level index is often used to estimate beech mast (Eichhorn, et al. 2016). It covers a range from ‘absence of fruits’ (0) up to ‘abundant fruits’ (3). In the study presented here, the beech mast index was used to calculate a factor by which the annual number of harvested wild boar was adjusted (for details see Section ‘Computation of relative wild boar abundance’). The beech mast index values for the consecutive years 2011-2017 were 3, 0, 2, 1, 0, 3, 0 (Nussbaumer, et al. 2016). Including the beech mast index in the computation of relative wild boar abundance was based on the assumption that in rich mast years wild boar are harder to hunt, because they visit hunters’ baiting sites less frequently (Bozzuto and Geisser 2019). *Baiting* refers to the practice of hunters putting out food to attract wild boar in locations where they are known to be frequent.

#### Probability of wild boar occurrence

An area-covering data grid with the probabilities of wild boar occurrence for all 37,738 sq km raster cells of Switzerland (water and glaciers were excluded) in summer was produced in previous work (Vargas-Amado, et al. 2020). This data grid was used in this study to divide the relative abundance values computed for different areas of wild boar occurrence among the individual grid cells (for details see Section ‘Computation of relative wild boar abundance’).

#### Forest cover

The forest cover of the National Forest Inventory (NFI) (Waser, et al. 2015), together with data about rest areas along motorways and outdoor piggeries, was used to identify the areas where a direct transmission of a disease from wild boar to domestic pigs is more likely.

#### Motorways and rest areas

The national routes were downloaded on September 8, 2020, from the Federal geoportal ‘geo.admin.ch’.^2^ The shapefiles of all 182 rest areas were obtained from the same source and from the Bundesamt für Landestopografie swisstopo along with the product swissTLM3D 2020.^3^

#### Agricultural zones boundaries

The agricultural zones boundaries, version from 2017, were downloaded from ‘geo.admin.ch’ in order to mark off areas for summer grazing of domestic pigs.^4^

#### Outdoor piggeries

Data about the geographical location and type (solid run area vs. pasture) of outdoor piggeries for the years 2011–2019 were obtained from the Federal Office for Agriculture (FOAG). The number of piggeries was not stable over the observation period. In 2019 there were 3,085 holdings in the RAUS program (‘Regelmässiger Auslauf im Freien’) with a solid run area (without pasture) and 344 holdings with pasture. The two types of outdoor piggeries were accurately described in a related publication (Früh 2011). In addition, the geographical locations of Alpine pastures, where pigs labeled as ‘Alpschwein’ graze in summer, were manually extracted from the map on the relevant web site.^5^ There is no comprehensive list of such pastures in Switzerland. The extracted ones are examples used to find out whether the dynamics of the husbandry system could be a driver of seasonal variation in transmission risk.

### Data analyses

Figure 1 shows the model of proposed ASF transmission with risk factors and model variables. The components of the model are described in Section ‘Computation of relative wild boar abundance’, ‘Estimation of the risk of disease introduction’, ‘Estimation of the risk of disease transmission’, and ‘Estimation of the combined risk of disease introduction and transmission’. Computation of the relative wild boar abundance is given some emphasis, because it refines the state of the art in a way not previously reported.

**Figure 1.**
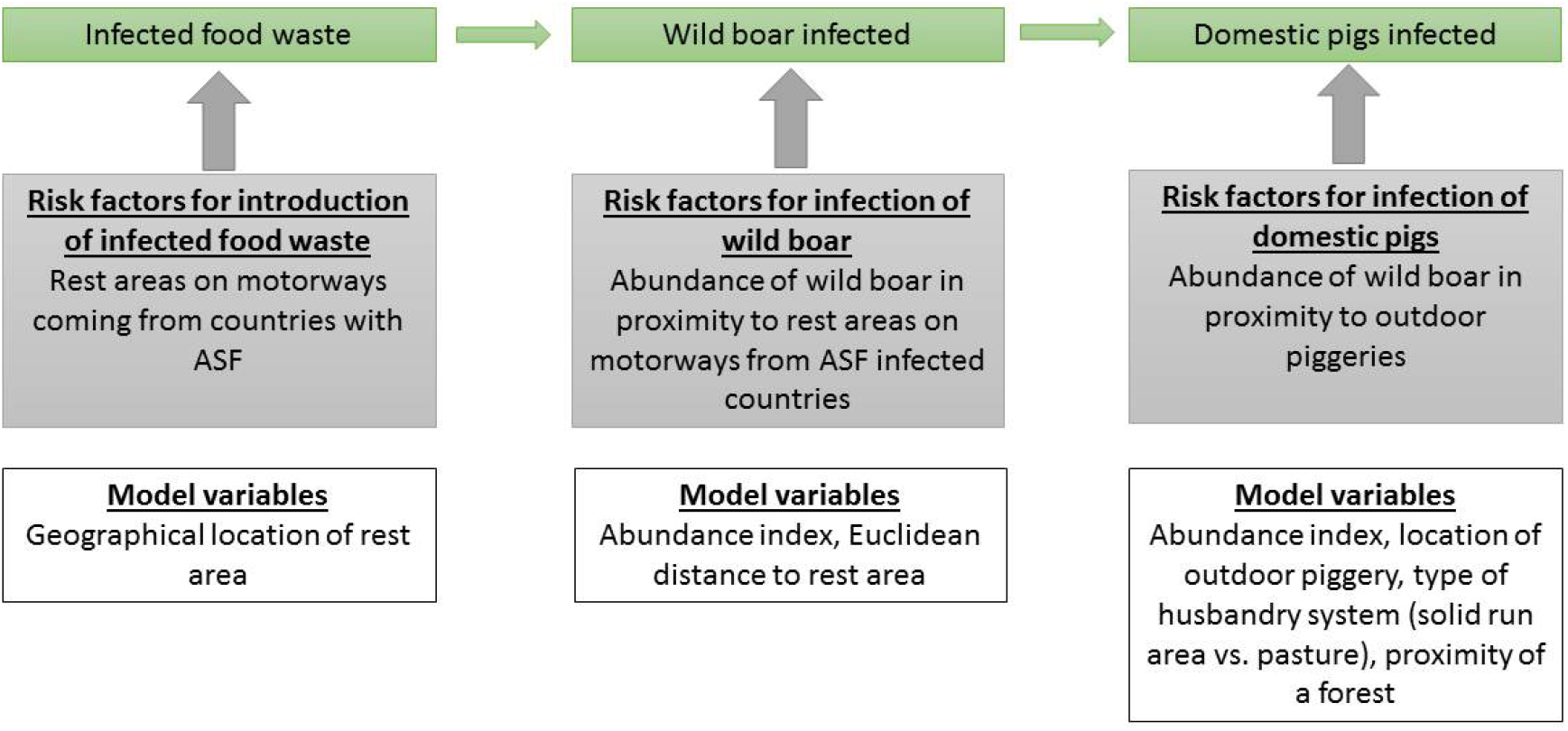
Model of proposed ASF transmission (green boxes) with risk factors (grey boxes) and model variables (white boxes)

#### Computation of relative wild boar abundance

For all cantons with wild boar occurrence, relative abundance was computed as an index value per sq km for summer (i.e., after reproduction and before hunting). *Relative abundance* refers to the “relative representation of a species in a particular ecosystem.” It reflects the “temporal or spatial variations of the size or density of a population but does not directly estimate these parameters” (ENETWILD-consortium, et al. 2018, 8) . In the work presented here, the *spatial* variations of the size or density of the wild boar population in Switzerland were of particular interest. The following equation expands on related work (ENETWILD-consortium, et al. 2018) by including additional factors relevant to relative wild boar abundance:

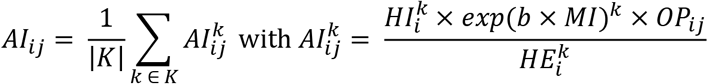

*AI_ij_* is the abundance index value of cell *j* in area *i* averaged over the observation period; the resulting real number was assigned to one of five index classes (‘not present’, ‘low’, ‘low–medium’, ‘medium–high’, ‘high’) based on the value range in which it fell.

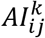 is the abundance index value of cell *j* in area *i* for hunting year *k*.

|*K*|is the number of hunting years; a hunting year is the period between March 1 to February 28 of the following year.

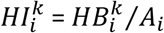 is the hunting index for hunting year *k* in area *i*.

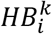is the hunting bag, i.e., the number of boars shot during hunting year *k* in area *i*. It is important to note that most Swiss cantons do not have quotas for wild boar; Neuchâtel has a quota which, according to the competent authority, has never been exploited; the canton of Jura has quotas for boars > 50 kg, but not for lighter ones

*A_i_* is the size of area *i* in square kilometers.

*exp*(*b* × *MI*)*^k^* is a factor adjusting the effect of mast conditions on hunting success in hunting year *k* (for details see below).

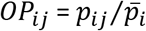is the (relative) probability of wild boar occurrence of cell *j* in area *i*.

The probability of wild boar occurrence of cell *j* in area *i* (i.e., *p_ij_*) was computed for the closed season for hunting in previous work using a number of statistical models of suitable wild boar habitat (Vargas-Amado, et al. 2020), 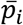 is the mean probability of wild boar occurrence of all cells in area *i*.

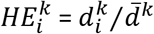 is the hunting effort for hunting year *k* in area *i* in terms of (relative) number of hunting days, 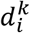is the number of hunting days in area *i* for hunting year *k*, *d* 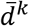is the number of hunting days for hunting year *k* averaged over all areas; the number of hunters hunting wild boar was not available in this study, nor was there sufficient information about the hunting method.

Area *i* was established based on the pooled hunting data for the entire observation period. Data were pooled to balance short-term variations in the spatial distribution of yearly harvested wild boar that were not assumed to be related to colonization/decolonization. How area *i* was established depended on the spatial granularity of the hunting data available in a canton. For cantons reporting mere counts per commune, hunting ground, or canton, these were the spatial units to which the equation was applied (see Table 1). When the data came with geographic coordinates, the commune, which is the lowest level of administrative division, or hunting ground in which a wild boar was shot was used as area *i*. Data with coordinates were handled this way in order to account for the animal’s ranging behavior. Overall, 1,004 areas were established.

The factor *b* × *MI* was proposed in a state-space model to estimate the (absolute) abundance of wild boar (Bozzuto and Geisser 2019). For a given hunting effort, *b* × *MI* is the rate by which the instantaneous harvesting mortality rate is adjusted based on mast conditions. Thereby, *MI* ∊ {0, 1, 2, 3} is the beech mast index and *b* = 0.023 is a scaling factor as estimated in the canton of Thurgau for the period of 1982–2017. Since beech mast in most years is a large-area phenomenon (Nussbaumer, et al. 2016), the same factor *b* was also used in the study presented here for other cantons with the same hunting system as Thurgau, namely Zurich, St. Gallen, Aargau, Solothurn, Basel-Stadt, Basel-Landschaft, Schaffhausen, and Luzern. For all other cantons, in which baited hunting is not practiced, the rate *b* × *MI* was set to 0. Given *b* × *MI*, the antilogarithm *exp*(*b* × *MI*) approximates the factor by which the hunting bag must be multiplied to account for mast conditions. It is important to note that this factor only balances the effect of mast on *hunting success*, which is a measure of how efficient hunting with a given effort is. The effect of mast on winter survival and reproduction is directly reflected in the hunting bags of the same and the following year. Figure 2 summarizes the workflow for the computation of relative abundance from hunting data.

**Figure 2.**
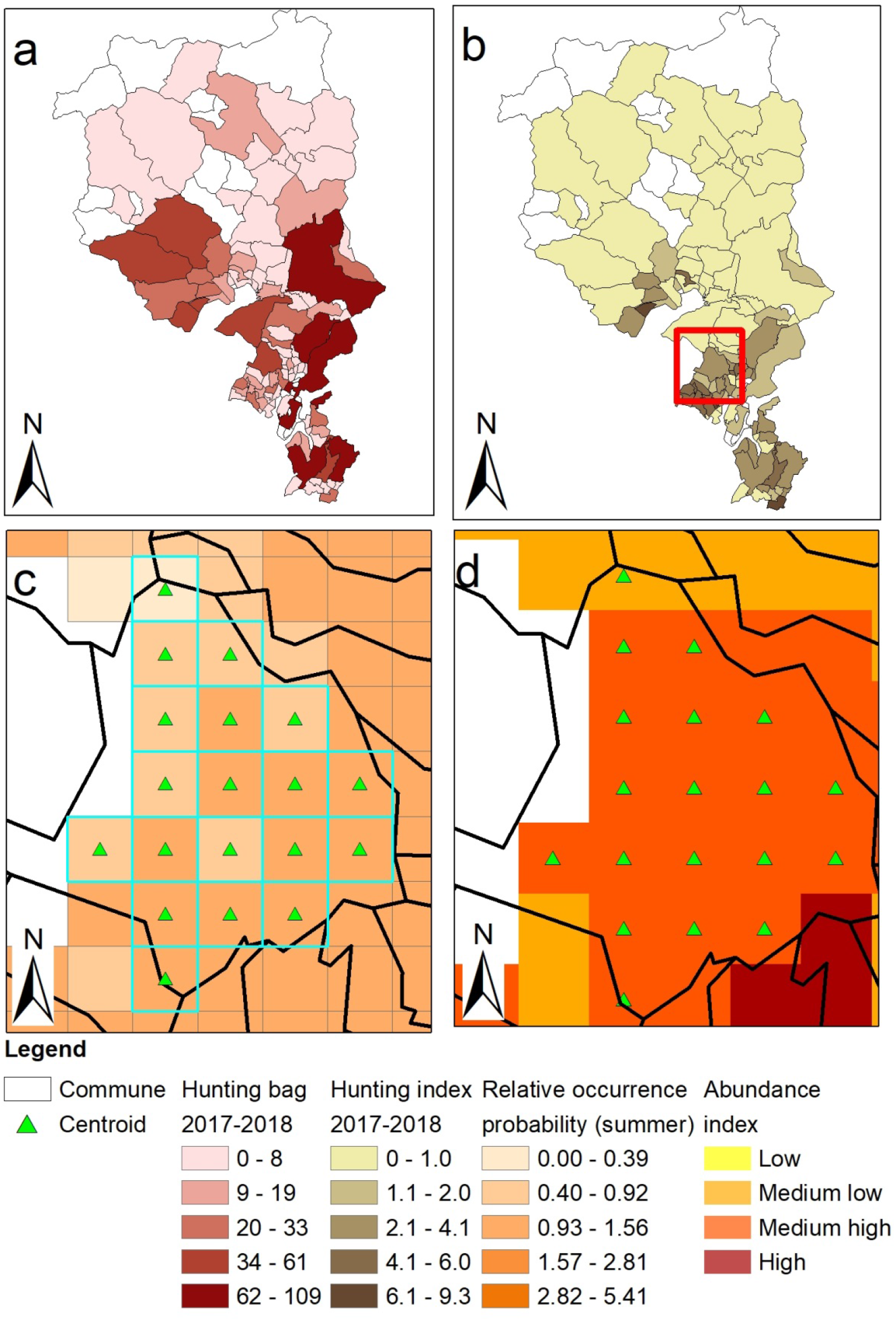
Step-by-step computation of the abundance index for hunting year 2017/18 in the canton of Ticino. a: hunting bag per commune, b: hunting index per commune, c: relative probability of wild boar occurrence in summer as a country-wide data grid (1 sq km), d: abundance index per grid cell.

#### Estimation of the risk of disease introduction

According to the National program for early detection of ASF (Bundesamt für Lebensmittelsicherheit und Veterinärwesen BLV 2020a), contaminated food waste that is discarded carelessly poses the highest risk of disease introduction into Switzerland. Rest areas along motorways in wooded areas are considered particularly exposed to this way of introduction, because motorways connect ASF affected countries to the urban centers and wooded areas are the preferred habitat of wild boar. Relevant motorways were identified by searching for the fastest routes from Bulgaria (Sofia), Hungary (Budapest), Romania (Bucharest, Cluj-Napoca, Timișoara, Iași), Poland (Warsaw, Kraków, Wrocław, Poznań, Gdańsk), Serbia (Belgrade), and Slovakia (Košice) to Switzerland (Zurich, Geneva, Basel, Bern, Lausanne) using Google Maps’ route planner and by looking up the main transit roads for heavy goods traffic through Switzerland on ‘map.geo.admin’. The points of departure were selected based on the map of the Friedrich-Loeffler-Institut (FLI), where all cases of ASF in Europe are cumulatively displayed for every calendar year.^6^ Routes were searched on October 7–8, 2020.

Table 2 shows the number of potentially exposed rest areas along the routes from 13 cities in ASF affected countries to five urban centers and along the main transit roads for heavy goods traffic through Switzerland per canton. Euclidean distance to the nearest rest area was calculated for each cell of a country-wide 1 sq km grid. Distances were classified into classes (1–4) to generate the scores for the calculation of the combined risk of disease introduction and transmission (see below).

**Table 2.**
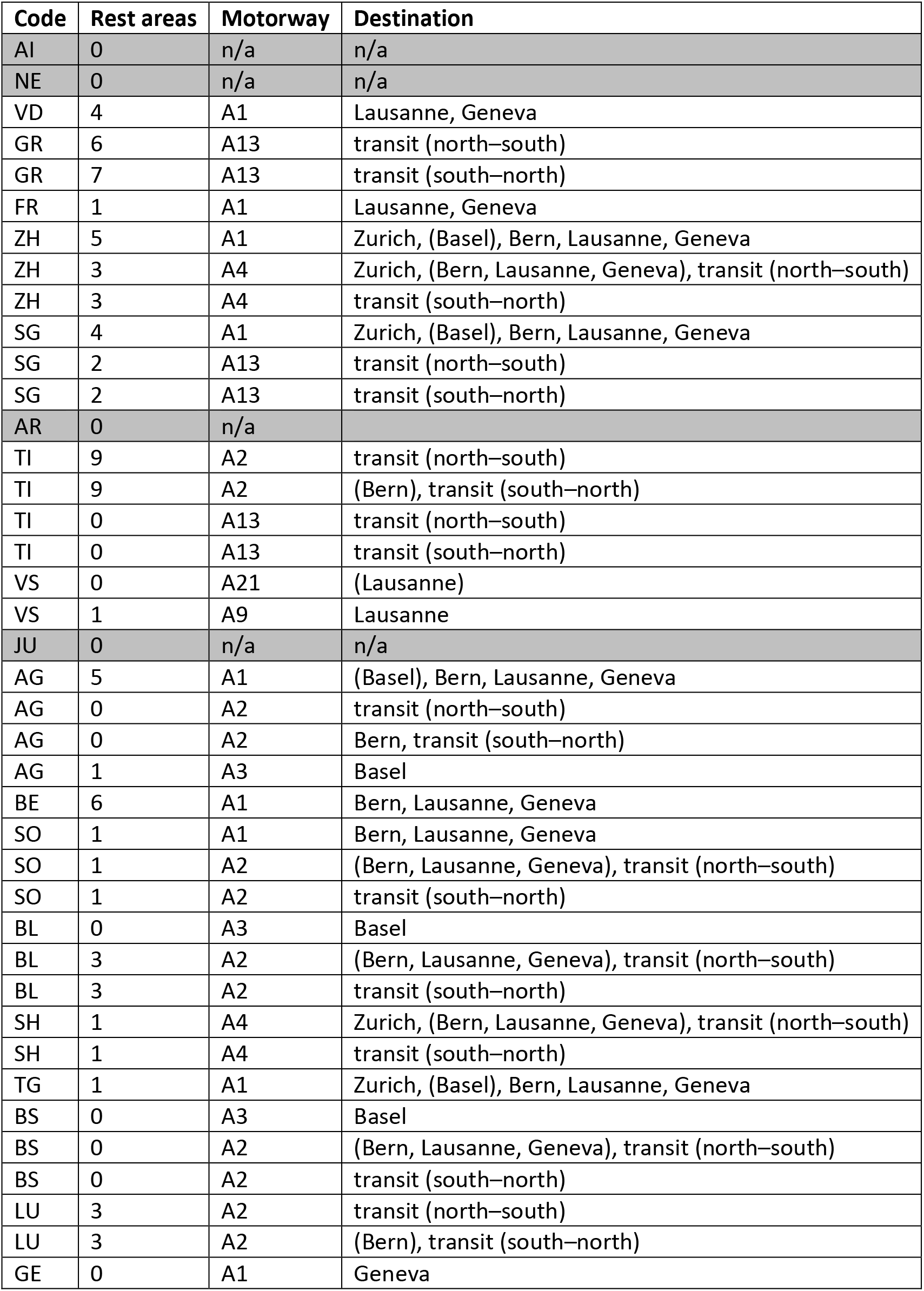
Number of potentially exposed rest areas along relevant motorways per Swiss canton (86 in all). Destinations in brackets indicate indirect connections. Motorways A1, A3, A9, A21 are traveled in one direction only; motorways A2, A4, A13 are traveled in both directions.

#### Estimation of the risk of disease transmission

Among the measures used for protecting domestic pig populations from a disease like ASF, the Federal Food Safety and Veterinary Office (FSVO) advocates not allowing pigs to contact with wild boar and, after an ASF outbreak, to avoid outdoor farming in areas infected by the disease (Bundesamt für Lebensmittelsicherheit und Veterinärwesen BLV 2020b). In order to identify risk areas for disease transmission to domestic pigs in the work presented here, communes with piggeries with a solid run area and communes with piggeries with pasture were located separately using the relevant toolset in ArcGIS.^7^ For each of the identified communes, piggery density was calculated by dividing the number of piggeries by the surface area of the commune. The resulting values were classified into classes (0–4) to generate the scores for the calculation of the combined risk of disease introduction and transmission and to ease the interpretation on the map.

#### Estimation of the combined risk of disease introduction and transmission

The combined risk of disease introduction and transmission reflects the risk of introducing a disease into the domestic pig population by the intermediary of wild boar. The combined risk was estimated by multiplying the values of relative wild boar abundance (scores 0–4), Euclidean distance to the nearest rest area (scores 1–4), density of outdoor piggeries (scores 0–4), and proximity of a forest (not shown). Proximity of a forest was assessed based on the forest cover NFI, where ‘wooded’ was given a score of 2, and a score of 1 was given otherwise. The values resulting from the multiplication were classified into classes ‘no risk’ (score 0), ‘low’ (scores 1–12), ‘medium low’ (scores 13–24), ‘medium high’ (scores 25– 48), ‘high’ (scores 49–128) based on the relative distribution of scores for piggeries with pasture. The resulting raster layer was transformed to a feature layer and the final scores were generalized to yield a single value per commune based on the maximum cell value in that commune. This was carried out in order to identify the political units in which risk areas were found and to facilitate the interpretation on the map. It was accomplished separately for piggeries with a solid run area and for piggeries with pasture. The consideration of proximity of a forest and type of husbandry system (solid run area vs. pasture) was motivated by a related study in which these were identified as risk factors for a contact between wild boar and outdoor pigs (Wu, Abril and Thomann, et al. 2012).

## Results

### Relative abundance of wild boar

Figure 3 shows the relative abundance of wild boar in Switzerland. The northern wild boar population ranges from Geneva to St. Gallen, covering most parts of the Jura and the adjacent regions of the Central Plateau, the Lower Valais, and the Lower Rhine valley. Wild boar occur occasionally also in the Upper Valais, the valleys of the Berner Oberland, and in the canton of Luzern (for the cantonal boundaries see Figure 4–6). This population is contiguous with the wild boar populations in neighboring Germany and France. The southern population is located in the canton of Ticino and in the region of Moesa in Graubünden, but is contiguous with the northern Italian wild boar population.

**Figure 3.**
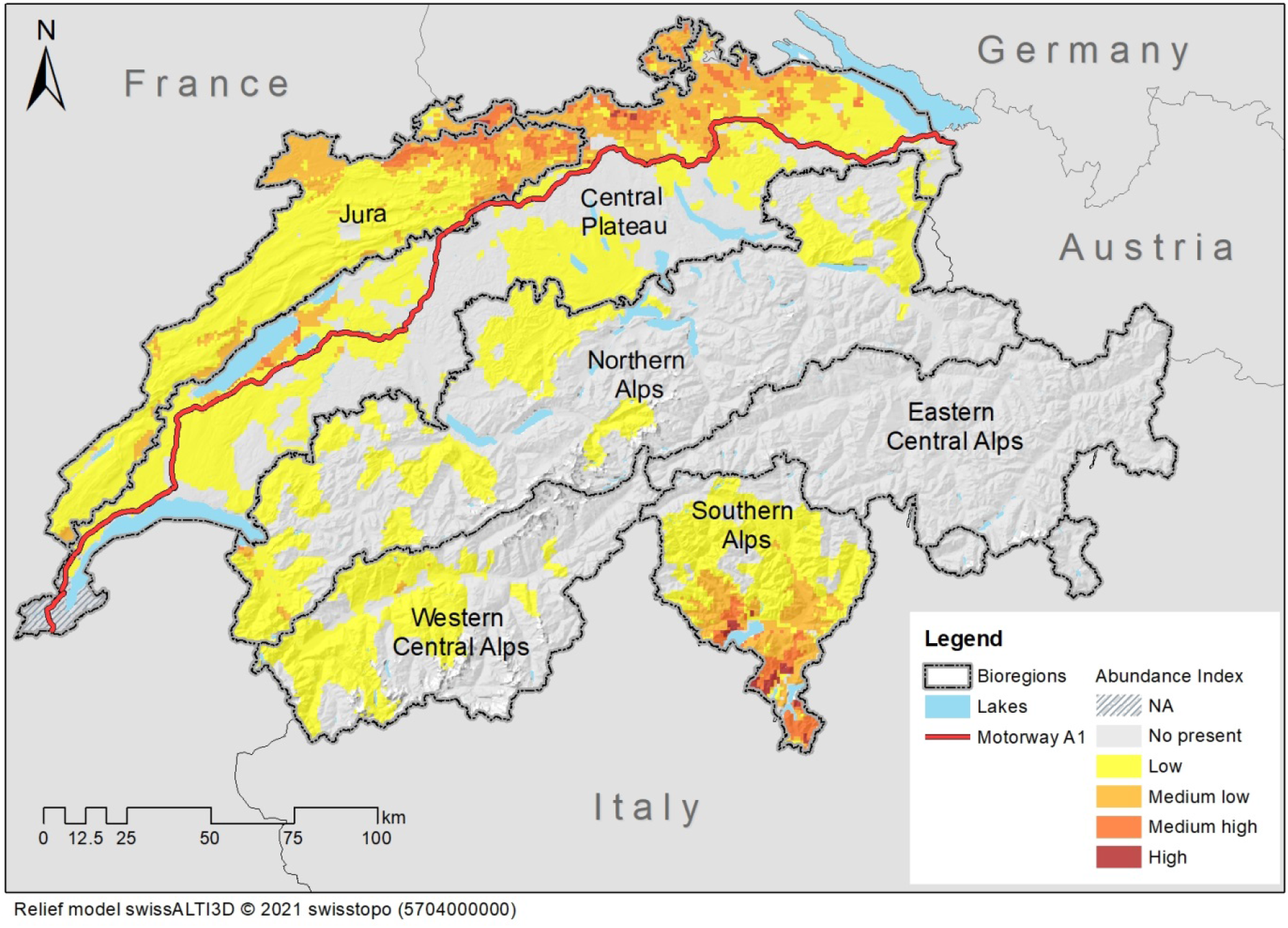
Relative abundance of wild boar in Switzerland. The numerical values underlying the nominal index values are not shown to avoid that these are mistaken as (absolute) wild boar ‘densities’.

**Figure 4.**
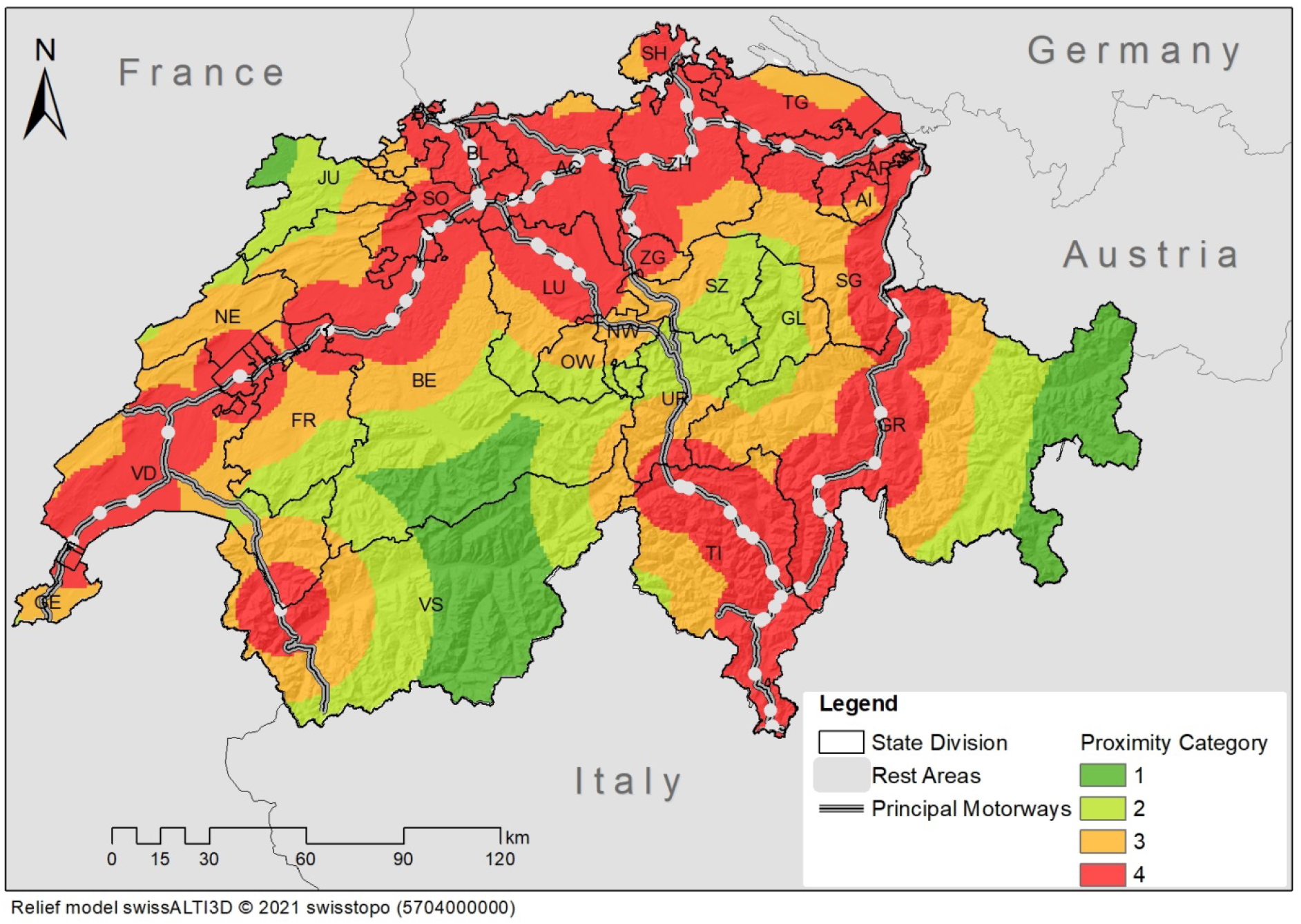
Euclidean distance to the nearest rest area along one of the routes from 13 cities in ASF affected countries to five urban centers in Switzerland and the main transit roads for heavy goods traffic through Switzerland. Routes were identified using Google Maps’ route planner (65 trips in all), they lead to motorways A1, A3, A9, and A21. The main transit roads for heavy goods traffic through Switzerland were motorways A2, A4, A13.

**Figure 5.**
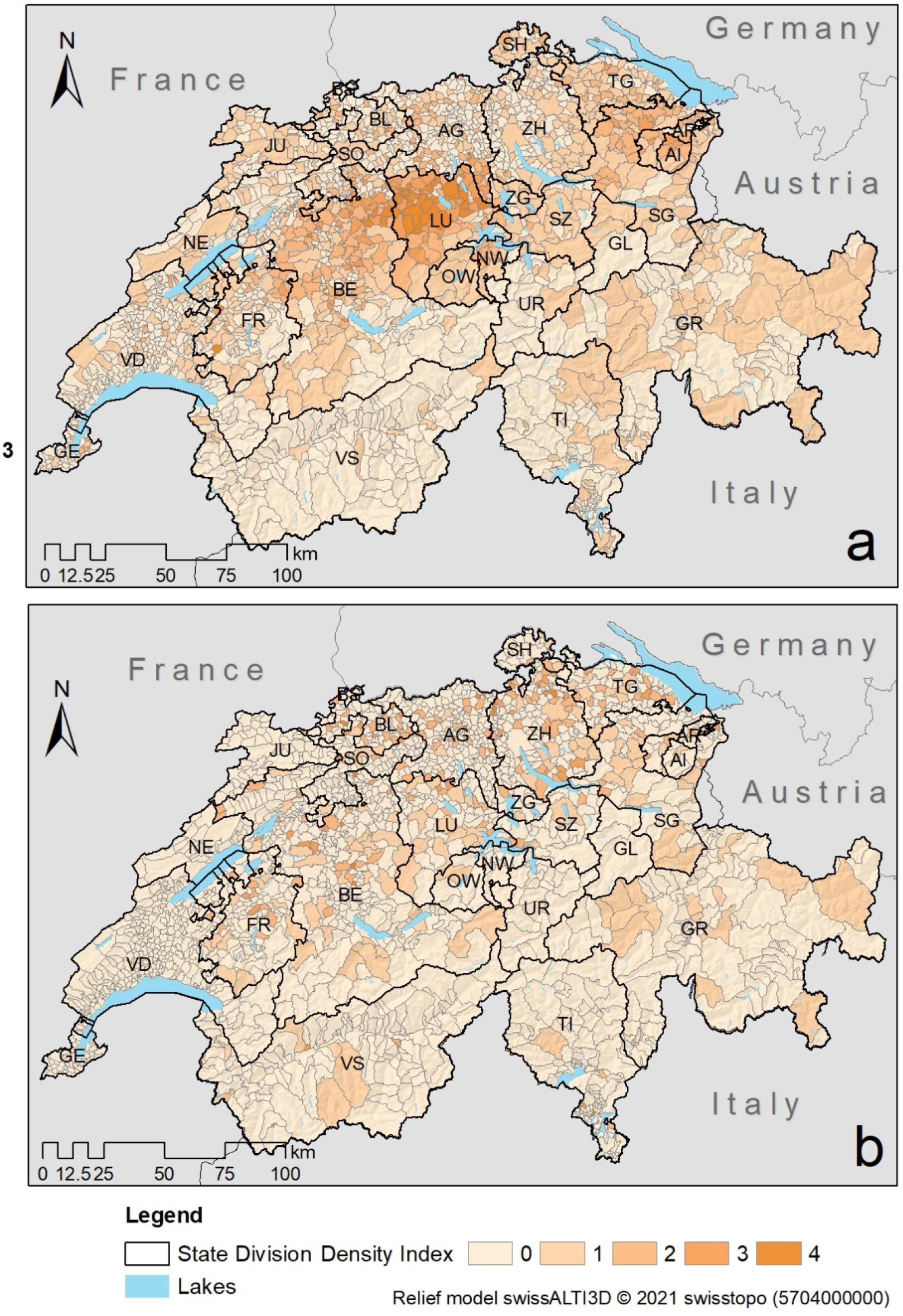
Density of outdoor piggeries. a: piggeries in the RAUS program (i.e., run area without pasture); RAUS stands for ‘Regelmässiger Auslauf im Freien’. b: piggeries with pasture.

**Figure 6.**
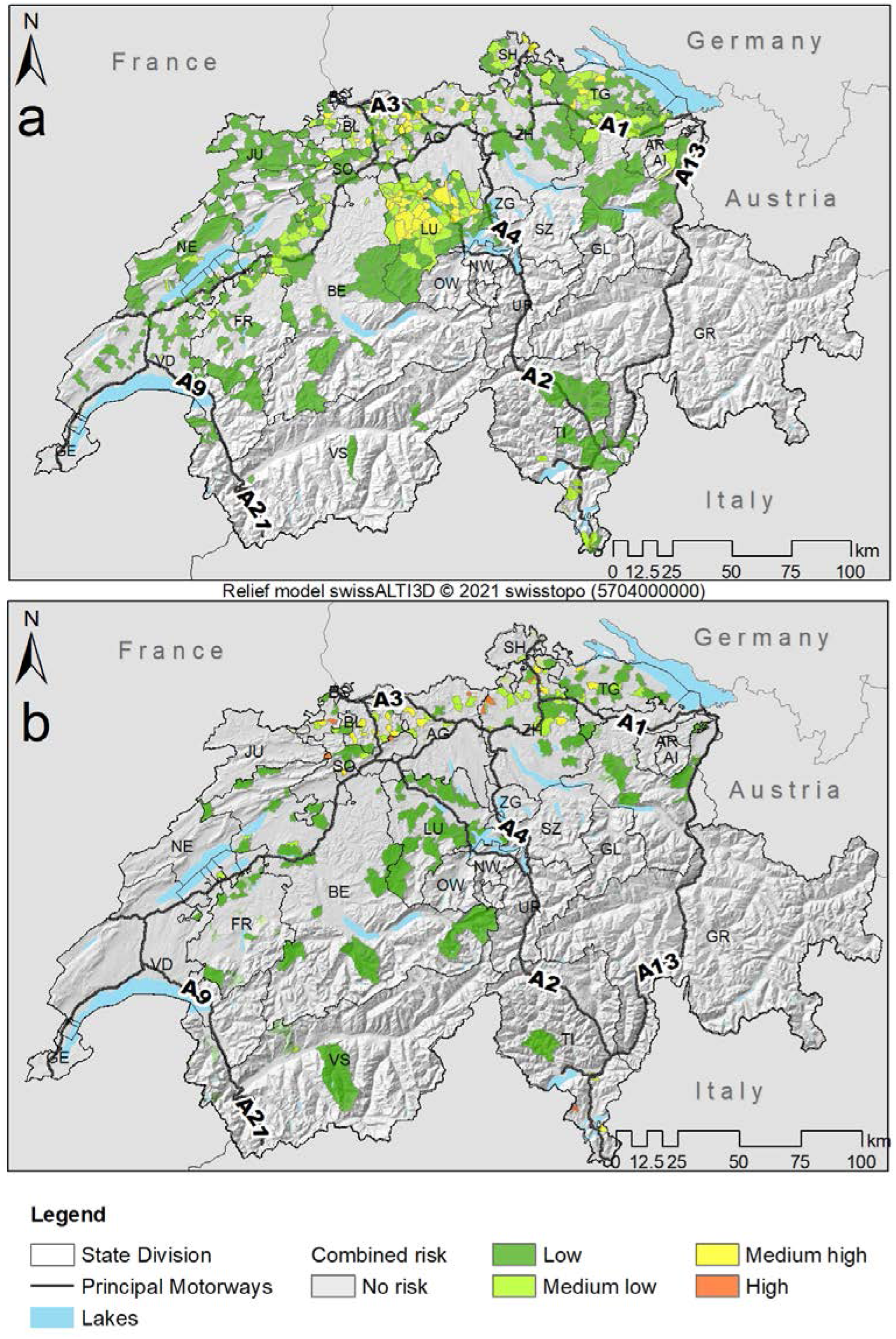
Areas with a combined risk of disease introduction into the wild boar population and transmission to domestic pigs identified based on wild boar abundance, Euclidean distance to the nearest rest area, density of outdoor piggeries, and proximity of a forest. a: piggeries in the RAUS program (i.e., run area without pasture). b: piggeries with pasture.

Wild boar are most abundant in areas near the borders of France, Germany, and Italy. They are also abundant in the south-east of Lake Neuchâtel. A number of reserves for waterbirds and migratory birds are located there, in which hunting is prohibited. In the Alpine canton of Ticino the spatial pattern of relative wild boar abundance is not only governed by the distance from the border, but also by the meters above sea level: wild boar range in areas above the tree line around 2,000 meters (not shown) only sporadically.

The spatial pattern of relative wild boar abundance suggests that motorway A1 is a barrier for wild boar colonizing Switzerland from the north in the canton of St. Gallen and parts of Thurgau. It is also a barrier for wild boar colonizing Switzerland from the north-west between Zurich and Bern. Motorway A1 is a leaky barrier between the rest area Hexentobel (TG) and Zurich as well as west of Bern.

### Risk areas for disease introduction

Figure 4 shows the proximity categories in which the cells of a country-wide data grid fall when classified according to the Euclidean distance to the nearest rest area along one of the relevant motorways. Ninety-six rest areas are not along motorways connecting ASF affected countries to Switzerland (not shown). Fifty-seven out of the displayed 86 rest areas are located in areas ranged by wild boar. These are the most likely hot spots for disease introduction into the Swiss wild boar population. They are listed by name below.

AG Walterswil, Würenlos
BE Lindenrain, Oberbipp-Nord
BL Mühlematt (both directions), Pratteln-Süd, Sonnenberg (both directions)
FR Rose de la Broye
GR Campagnola (both directions)
LU Chilchbüel, Inseli, Knutwil-Nord, Knutwil-Süd, Neuenkirch (both directions)
SG Rheintal Ost, Rheintal West, Thurau Nord, Wildhus Nord
SH Berg, Moos
SO Eggberg, Gunzgen-Nord, Teufengraben
TG Hexentobel Nord
TI Bellinzona Nord, Bellinzona Sud, Bodio, Coldrerio (both directions), Giornico, Lavorgo (both directions), Moleno Nord, Moleno Sud, Motto, Muzzano (both directions), San Gottardo-Sud, Sasso, Segoma (both directions)
VD Bavois, Crans-près-Céligny, St-Prex
VS Dents de Morcles
ZH Baltenswil-Nord, Büsisee, Chrüzstrass, Forrenberg Nord, Kemptthal, Stegen, Weinland (both directions)

### Risk areas for disease transmission

Figure 5 shows the densities of outdoor piggeries at the level of communes. The spatial distribution of communes with piggeries with a solid run area is the same as that of communes with all types of piggeries (Sterchi, et al. 2019). High densities of piggeries with a solid run area are present in the cantons of Bern, Luzern, St. Gallen, Appenzell Innerrhoden, and Appenzell Ausserrhoden. By contrast, outdoor piggeries with pasture are more evenly distributed across Switzerland. Densities of piggeries were in the same range for both types of husbandry system, namely 0.004–1.880 piggeries per sq km and 0.004–1.167 piggeries per sq km, respectively. However, the mean was more than twice as high for piggeries with a solid run area than for piggeries with pasture (0.279 vs. 0.114). Accordingly, the fraction of communes with a low density is higher for piggeries with pasture, which is in line with the observation in Figure 5 that communes with extensive pig farming are not geographically connected.

### Areas with a combined risk of disease introduction and transmission

Figure 6 shows areas with a combined risk of disease introduction into the wild boar population and transmission to domestic pigs. Accordingly, domestic pigs are most at risk of becoming infected in outdoor piggeries located near the borders of France, Germany, and Italy. On the north side of the Alps, high risk areas are located north of the A1, the main motorway crossing the Central Plateau. Piggeries with a solid run area and piggeries with pasture differ in the size of the risk areas and particularly in the canton of Luzern also in their score.

Figure 7 shows examples of Alpine pastures within or in close proximity to areas ranged by wild boar, where pigs labeled as ‘Alpschwein’ graze in summer. Pigs are held in these areas in order to use some of the by-products of summer alpine cheesemaking. These pastures are not included in Figure 5, because a comprehensive list was not available. When compared with Figure 4, Figure 7 shows that some pastures in the cantons of St. Gallen and Ticino are located in the proximity of rest areas. The combined risk of disease introduction and transmission is low for all other pastures.

**Figure 7.**
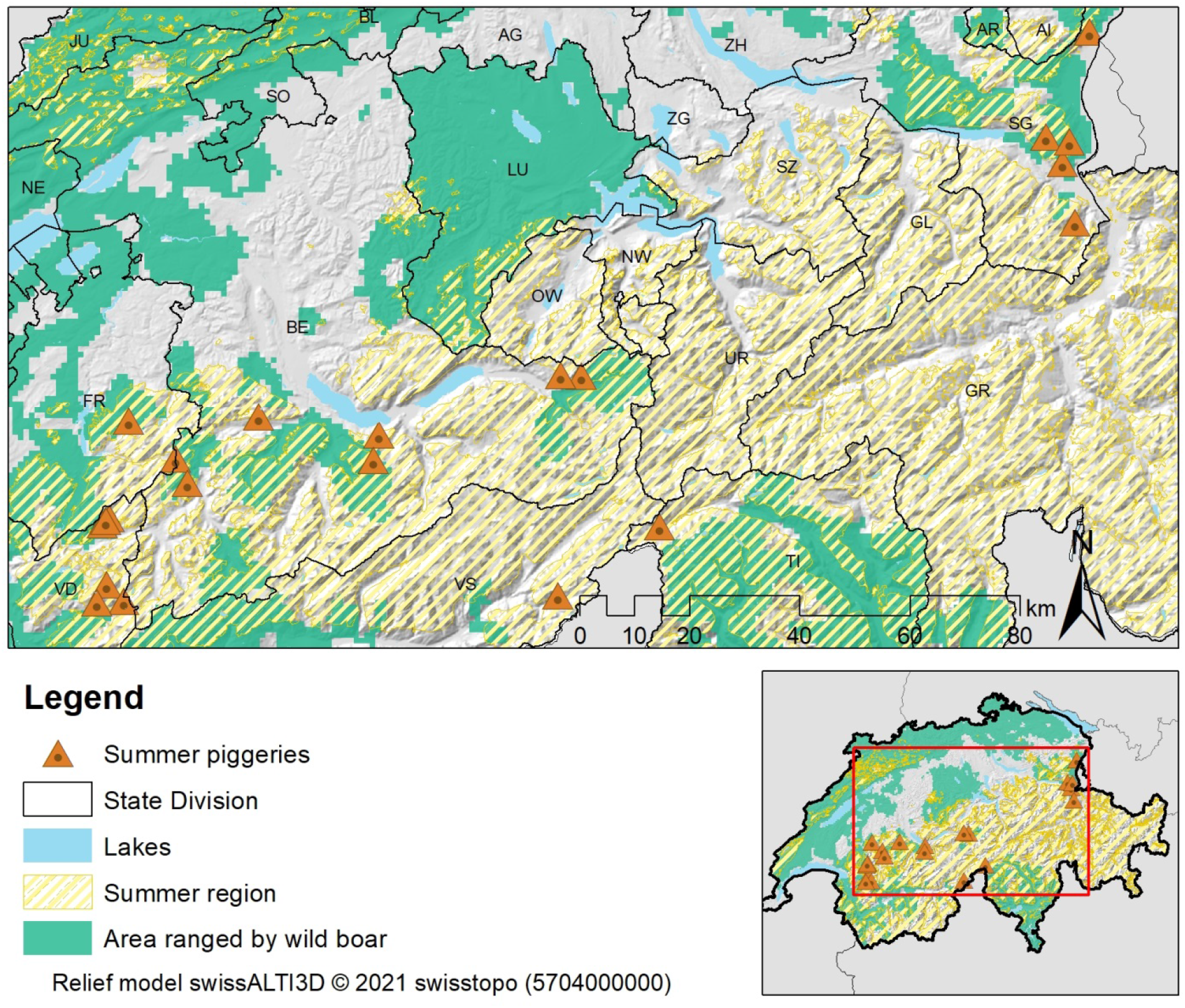
Alpine pastures, within or in close proximity to areas ranged by wild boar, where pigs labeled as ‘Alpschwein’ are grazed during summer.

## Discussion

A novel method was presented to estimate and map the risk of introducing ASF into the domestic pig population through wild boar intermediate hosts. It makes use of data about hunted wild boar, rest areas along motorways connecting ASF affected countries to Switzerland, outdoor piggeries, and forest cover. These data were used to compute relative wild boar abundance as well as to estimate the risk of both disease introduction into the wild boar population and disease transmission to domestic pigs. The way relative wild boar abundance was calculated adds to the current state of the art by considering the effect of beech mast on hunting success and the probability of wild boar occurrence when distributing relative abundance values among individual grid cells. The risk of ASF introduction into the domestic pig population by wild boar was highest near the borders of France, Germany, and Italy. On the north side of the Alps areas of high risk were located on the unshielded side of the main motorway crossing the Central Plateau, which acts as a barrier for wild boar.

The results of this study can be used to focus surveillance efforts for early disease detection on areas where the combined risk of disease introduction into the wild boar population and disease transmission to domestic pigs is high. African Swine Fever is currently at the center of attention in western European countries. However, the results of the analyses carried out in this study may also inform policies to control other diseases that are transmitted by direct contact from wild boar to domestic pigs. Depending on the transmission route, the results allow for a subtle differentiation. Domestic pigs in both types of outdoor piggeries may be exposed to infectious agents transmitted by aerosols such as *Mycoplasma hyopneumoniae*. A study based on genotyping of *M. hyopneumoniae* isolated from pig lungs during enzootic pneumonia outbreaks and lungs from wild boar in the close proximity to the affected pig farms confirmed transmission of the pathogen between domestic pigs and wild boar (Kuhnert and Overesch 2014). By contrast, spill-over of pathogen such as *Brucella suis* that are sexually transmitted is less likely in piggeries with solid run area than in piggeries with pasture. In a study of the risk factors for contact between wild boar and outdoor pigs in Switzerland, mating events were reported for holdings with pure pasture or mixed run-out only (Wu, Abril and Thomann, et al. 2012).

Direct contact is not the only way how ASF can be transmitted between wild boar and domestic pigs. In the sequel of the Belgian outbreak in 2018–2019 a panel of 34 national and international experts assessed the risk associated with different transmission routes semi-quantitatively (Mauroy, et al. 2021). Among 25 routes for ASF transmission from wild boar to domestic pigs, the experts considered ‘farmer’, ‘bedding material’, ‘veterinarian’, ‘professionals from the pig sector’, and ‘swill feeding’ most important in the Belgian epidemiological context. ‘Living wild boar’ together with ‘contaminated vegetal products (feed)’ and ‘hunter’ ranked sixth. This suggests that the ‘human factor’, which is considered in the study presented here for disease *introduction*, could potentially play a role also in disease *transmission* in Switzerland.

The barrier effect of motorway A1, observed in Figure 3, emphasizes the need to account for landscape configuration and fragmentation when assessing the effect of management regimes on the ranging behavior of wild boar (Fattebert, et al. 2017). More fine-grained landscape configuration and fragmentation should be considered when the results of this study are used at the local level. This is an important component of the strategy of the Federal Food Safety and Veterinary Office (FSVO) for reducing the risk of an introduction of ASF at a cantonal level (Bundesamt für Lebensmittelsicherheit und Veterinärwesen BLV 2019a, Bundesamt für Lebensmittelsicherheit und Veterinärwesen BLV 2019b). The results of the study presented here can be used to inform the cantons about which small scale geographical areas to concentrate their efforts for early disease detection and control.

As stated in Section ‘Relative abundance of wild boar’, Swiss wild boar populations are contiguous with those in France, Germany, and Italy. Therefore, a disease like ASF could also be introduced by improper disposal of contaminated food waste in a foreign rest area near the Swiss border. Figure 4 shows that the zones bordering potential risk areas in Germany and Italy already have a high score of 3 or 4. Accordingly, considering rest areas in Germany or Italy would not have a significant effect on the areas with a high combined risk of disease introduction and transmission identified in Figure 6. Things are different in neighboring France where motorway A36 from Beaune to Mulhouse passes close by the canton of Jura with a low score of 1–2 in Figure 4. The consideration of French rest areas could potentially increase the combined risk reported in Figure 6. Another important potential way of ASF introduction into the Swiss wild boar population is via hunting tourism. Hunters should be properly informed of the associated risks and appropriate biosecurity methods by the competent authorities.

There is no viable wild boar population in the canton of Luzern. The canton is ranged by a few dispersed animals only. Nevertheless, the risk of introducing ASF into the domestic pig population by wild boar was estimated to be medium (Figure 6). This is primarily due to Luzern’s practice of reporting hunting data as an aggregate for the entire canton (see Table 1), resulting in a positive score also in areas where there are no wild boar. Overestimating the risk of disease introduction in this canton does not have an adverse effect on the recommendations for action derived from Figure 6. The probability of a wild boar encounter is expected to increase in the future: wildlife corridors crossing important motorways, including A1 and A2, that were formerly interrupted are currently repaired and new corridors are being built to increase habitat connectivity.

It would be interesting to estimate the changing risk of disease transmission at different times of the year in a future study. This would require that temporal (or seasonal) data on wild boar abundance were available, which is currently not the case. The abundance data in this study were estimated only for the summer. Provided there are no quotas, the hunting bag, in the long run, is proportional to the size of the population before the hunting season starts (ENETWILD-consortium, et al. 2018). In this study, data were averaged over many years to balance out strong effects of non-controllable factors, such as weather conditions, on the annual number of harvested wild boar. Dealing with relative summer abundance does not limit the scope of this research. Summer is the season where the risk of transmission is highest for a number of reasons. First, the wild boar population is most abundant in summer after spring reproduction and before hunting. Second, the area potentially ranged by wild boar is larger in summer than in winter (Vargas-Amado, et al. 2020). Third, domestic pigs are grazed on Alpine pastures in summer. The seasonal variation in transmission risk may primarily be driven by the dynamics of the husbandry system, rather than by variations in wild boar abundance. Disease control agencies are well-advised to keep country-wide records of Alpine pastures with domestic pigs in the future.

In future, the degree of connectedness of piggeries to the rest of the domestic pig production network could be added in order to assess the consequences of a disease introduction. Such an extension should expand on previous work investigating the structure and patterns of the pig transport network in Switzerland (Sterchi, et al. 2019).

## Conclusions

This study has shown that risk-based surveillance for early detection of disease epidemics can benefit from approaches used in wildlife research and geographic information science. Combining these methods is especially advantageous when wildlife reservoirs are important for disease transmission, as the data that are needed for risk estimation are highly variable. Data preprocessing methods from either of these fields may be useful to prepare these data for analysis. Carrying out the analysis may also require techniques originating from wildlife research or geographic information science. In the study presented here, almost all variables had a spatial reference and the way they were modeled required some form of geographic information processing. Involving multiple disciplines is essential for providing the skills and methods needed to deal with the challenges posed by a disease emergence at the livestock-wildlife interface.

## Conflict of interest statement

None declared.

## Acknowledgements

The authors would like to thank Dominique Suter of the Federal Food Safety and Veterinary Office FSVO for providing references and commenting on the draft manuscript. They acknowledge the assistance of Barbara Früh of the Research Institute of Organic Agriculture FiBL and Félix Gréverath of the Federal Office for Agriculture FOAG with collecting and interpreting data about pig husbandry in Switzerland.

## Funding

This work was supported by the Swiss National Science Foundation (SNSF) NRP75 [grant number 407540_167303]. Maria J. Santos is funded by the University Research Priority Program in Global Change and Biodiversity.

## Footnotes

1 https://www.jagdstatistik.ch

2 http://map.geo.admin.ch/?layers=ch.astra.nationalstrassenachsen

3 https://www.swisstopo.admin.ch/en/geodata/landscape/tlm3d.html

4 http://map.geo.admin.ch/?layers=ch.blw.landwirtschaftliche-zonengrenzen

5 http://www.alpschweine.ch/

6 https://www.fli.de/de/aktuelles/tierseuchengeschehen/afrikanische-schweinepest/karten-zur-afrikanischen-schweinepest/

7 https://desktop.arcgis.com

## Notes

### Competing Interest Statement

The authors have declared no competing interest.

